# ModEx: A text mining system for extracting mode of regulation of Transcription Factor-gene regulatory interaction

**DOI:** 10.1101/672725

**Authors:** Saman Farahmand, Todd Riley, Kourosh Zarringhalam

## Abstract

**Background:** Transcription factors (TFs) are proteins that are fundamental to transcription and regulation of gene expression. Each TF may regulate multiple genes and each gene may be regulated by multiple TFs. TFs can act as either activator or repressor of gene expression. This complex network of interactions between TFs and genes underlies many developmental and biological processes and is implicated in several human diseases such as cancer. Hence deciphering the network of TF-gene interactions with information on mode of regulation (activation vs. repression) is an important step toward understanding the regulatory pathways that underlie complex traits. There are many experimental, computational, and manually curated databases of TF-gene interactions. In particular, high-throughput ChIP-Seq datasets provide a large-scale map or transcriptional regulatory interactions. However, these interactions are not annotated with information on context and mode of regulation. Such information is crucial to gain a global picture of gene regulatory mechanisms and can aid in developing machine learning models for applications such as biomarker discovery, prediction of response to therapy, and precision medicine.

**Methods:** In this work, we introduce a text-mining system to annotate ChIP-Seq derived interaction with such meta data through mining PubMed articles. We evaluate the performance of our system using gold standard small scale manually curated databases.

**Results:** Our results show that the method is able to accurately extract mode of regulation with F-score 0.77 on TRRUST curated interaction and F-score 0.96 on intersection of TRUSST and ChIP-network. We provide a HTTP REST API for our code to facilitate usage.

**Availibility:** Source code and datasets are available for download on GitHub: https://github.com/samanfrm/modex HTTP REST API: https://watson.math.umb.edu/modex/[type query]

## 1. Introduction

Gene regulatory networks are essential in many cellular processes, including metabolism, signal transduction, development, and cell fate [1]. At the transcriptional level, regulation of genes is orchestrated by concerted action between Transcription Factors (TFs), histone modifiers, and distal cis-regulatory elements to finely tune and modulate expression of genes. Sequence-specific TFs play a key role in regulating gene transcription at the transcriptional level. They bind specific DNA motifs to regulate promoter activity and either enhance (activate) or repress (inhibit) expression of the genes. Deciphering transcriptional regulatory networks is crucial for understanding cellular mechanisms and response at a molecular level and can shed light on molecular basis of complex human diseases [2, 3, 4, 5]. Moreover, knowledge on interactions between genes and biomolecules is an essential building block in several pathway inference and gene enrichment analysis methods that aim to annotate an altered set of transcripts with biological function [6].

A high-throughput experimental approach for identifying regulatory interaction is chromatin immunoprecipitation followed by sequencing (ChIP-Seq). In ChIP-Seq methodologies, antibodies that recognizes a specific TF are used to pull down attached DNA for sequencing. The ENCODE (Encyclopedia of DNA Elements) consortium [7] has produced vast amount of publicly available high-throughput ChIP-Seq experiments that are processed and deposited into databases such as GTRD [8] and ChIP-Atlas [9] (>40,000 human experiments). These databases can be utilized to construct a high coverage transcriptional regulatory network. There are also other sources of transcriptional regulatory network including JASPAR [10], the Open Regulatory Annotation database (ORegAnno) [11], SwissRegulon [12], the Transcriptional Regulatory Element Database (TRED) [13], the Transcription Regulatory Regions Database (TRRD) [14], TFactS [15], TRRUST [16]. These databases have been assembled with a variety of approaches, including reverse engineering approaches based on high-throughput gene expression experiments [17, 18], text mining approaches [19], and manual curation [20].

Although these databases are a valuable source of gene regulatory information, there are several constraints that limit their usability. For instance, databases of computationally predicted and expression-driven interactions are typically very noisy. Importantly, the majority of the databases including ChIP-derived databases do not report the mode of regulation (up or down) – which is crucial to understanding the functional behavior of the cell. In this study, we propose a text mining system ModEx, to mine biomedical literature and annotate ChIP-derived regulatory interactions.

The rest of the paper is structured as follows. In Section 2, we briefly review some backgrounds and related works. In Section 3, we describe the datasets and present the details of the proposed information extraction and event extraction components and introduce our regulatory mode extraction using a long-range dependency graph. System evaluation and benchmark results are presented in Section 4. We conclude the paper and discuss the result, limitations, and future work in Section 5.

## 2. Overview and related works

Text mining plays an important role in unveiling purified information from a large number of documents in a satisfactory time. Essential steps for biomedical text mining can be divided into 3 steps: (1) information retrieval (IR), (2) name entity recognition (NER), and (3) information extraction (IE). Together, they can be utilized to identify specific biological knowledge from literature [21, 22].

IR tools retrieve relevant text information from articles, abstracts, paragraphs, and sentences corresponding to subject of interest. A popular IR approach for biomedical application is the use of Boolean model logic (AND/OR) for extracting relevant information containing specific biological terms [23]. Prominent IR tools that use the Boolean logic model are iHOP [24] and PubMed. PubMed utilizes human-indexed MeSH terms to reduce the search space and retrieve relevant abstracts containing user specified keywords. iHOP builds on PubMed and is able to detect co-occurrence of terms. A limitation of iHOP is that the terms must occur in the same sentence.

After the IR step, NER must be used to identify relation between biological entities. This is a challenging step as entity names are not unique. Therefore, NER tools must take textual context into consideration to accurately detect entities. For example, gene names may have different variations in orthographical structure (e.g. ABL1, Abl1, Abl-1) or multiple synonyms (e.g. ABL1, ABL, CHDSKM, Abelson tyrosine-protein kinase 1). ER methods, typically divide the task into two steps, (1) identify the entities and their location in the context, and (2) assign unique identifiers to the entities [23]. Fortunately, multiple terminological databases, such as Gene Ontology [25], UMBLS [26], BioLexicon [26], and Biothesaurus [26] provide information on biological entities and name variations and can be used to detect biological entities such as genes or proteins [27, 28].

Lastly, Relation Extraction (RE) is a task for extracting pre-defined facts relating to an entity or entities in the text [29]. In biomedical domain, multiple RE methods have been developed to extract information relating to genes [16], such as Mutation-Disease associations, protein-protein interaction [30, 31], pathway curation [32], gene methylation and cancer relation [33], biomolecular events [34], metabolic reactions [35] and gene-gene interactions [36]. For gene regulatory networks, which is the focus of this paper, the RE system must detect and extract a causal relation between a protein and a gene (e.g., A regulated B). This task is very complex, even for human experts [37]. To illustrate, consider the causal relation *“aatf upregulates c-myc”* that should be deduced from the following sentence: *“down-regulation of c-myc gene was accompanied by decreased expressions of c-myc effector genes coding for htert, bcl-2, and aatf”* [38]. Extracting a positive regulatory interaction between AATF and c-Myc is quite challenging using simple RE methods. For example, the RE method, may naively annotated the interaction as negative because of the keyword *“decreased”*. However, by taking *“down-regulation”* into account, the RE method would able to correctly extract a positive regulation from this sentence.

Therefore, construction of a causal transcriptional regulatory network by traditional means of text mining is hampered by these challenges and as a result, fully automated text-mining based models are limited in their scope and accuracy [23]. Combining experimentally derived regulatory interactions from high-throughput sources with text-mining approaches can bridge the gap between the two approaches and address their shortcomings.

In this work, we present a hybrid model ModEx, to mine the biomedical literature in MEDLINE to extract and annotate causal transcriptional regulatory interactions derived from high-throughput ChIP-seq datasets. Our model incorporates three main components of IR, NER and IE customized for mining regulatory interactions. Several expert-generated dictionaries are provided to optimize and complement the IR component. We proposed a weighted long-range dependency graph to extract causal relations and annotated the retrieved interaction with meta-data, such as full supporting sentences, PubMed ID, and importantly mode of regulation. Our pipeline bypasses several of the challenges of fully automated text-mining methods, including query translation for a particular interaction, relevant citation retrieval, entities recognition and regulatory annotation. ModEx was able to achieve an F-score 0.76 in retrieving and annotating a gold-standard regulatory network. We also compared ModEx with a state-of-the-art method, and the result shows strong improvement in terms of classification metrics.

## 3. Materials and methods

### 3.1. Datasets

We obtained TF-gene interaction data from ChIP-Seq experiments, deposited on the ChIP-Atlas database [9]. ChIP-Atlas contains all publicly available high-throughput ChIP-Seq experiments. We assembled regulatory networks from these interactions using various cutoff criteria for ChIP-Seq peak signal score and distance to the TSS. The least stringent criterion results in a network with 4 million interactions between 758 TFs and 18,874 target genes. However, there is no reported mode of regulation in ChIP-Atlas.

We used PubMed engine to query the MEDLINE database using the entities involved in interaction in ChIP-Atlas. MEDLINE is openly accessible and provides more than 25 million biomedical and life sciences references from approximately 5,600 worldwide journals. PubMed takes a query including keywords from user, and returns a list of citations that match input query.

Finally, TRRUST regulatory network [20] was utilized as gold standard to evaluate the performance of ModEx. TRRUST is a manually curated database of human transcriptional regulatory network with partial information on mode of regulation. It contains 9,396 regulatory interactions of 800 human transcription factors, 5,066 of which are annotated with information on mode of regulation (3,148 repression and 1,918 activation).

### 3.2. Information retrieval module

We developed an IR module, using Biopython [39], to retrieve the information from the MEDLINE for regulatory interactions in ChIP-Atlas. Figure 1 illustrates the overall workflow of our IR module to fetch relevant citations associated with the regulatory interaction. We start by building a query based on the entities participated in the interactions to retrieve abstracts from PubMed engine. PubMed engine takes free-text keywords and returns a list of ranked citations that match input keywords. Its search strategy has two major characteristics: first, it adds Boolean operators into user’s query and then uses automatic term mapping (ATM) [40].

**Figure 1:**
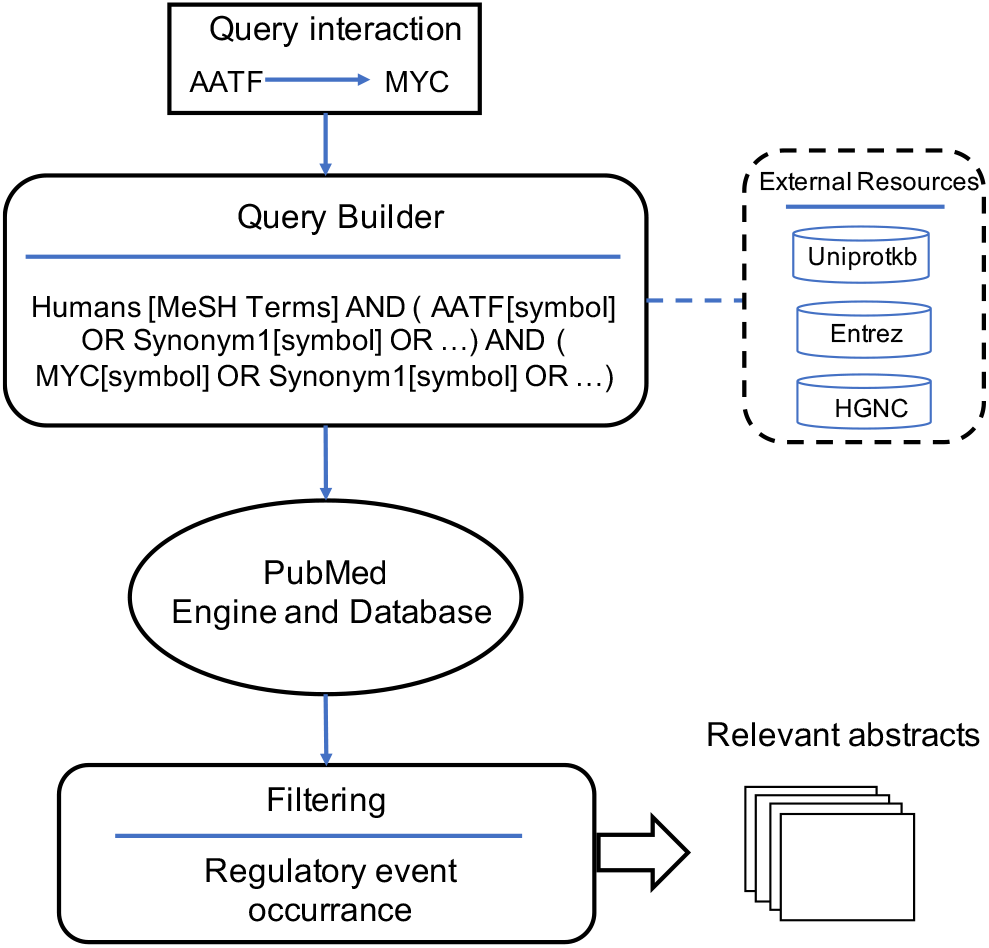
The Information Retrieval workflow. The steps are as follows: first, a Boolean query is built according to the associated entities in the regulatory interaction. It uses our local dictionary integrated from several external databases to complement the query with more synonyms and aliases. Then, the query is submitted to the PubMed engine and abstracts are retrieved for processing. Abstracts with no regulatory events are excluded for further analysis.

Each query was supplemented with extra terms acquired from several external resources, including HGNC, Entrez, and UniprotKB to fetch more relevant abstracts. We integrated these synonyms into a local dictionary covering gene symbol, synonyms and official full name. A Boolean query was then created to enforce our own search logic to the ATM in order to increase the chance of attaining relevant citations and also to reduce the response time. The query was made with appropriate Boolean logic (AND/OR) on entities and their extra terms using the lookup dictionary. A MeSH descriptor term (e.g. Humans) was also incorporated in the query to further boost the mapping process on PubMed engine. For examples the query for AATF and MYC regulatory interaction is, “humans [msh] AND (AATF[sym] OR BFR2[sym] OR CHE-1[sym] OR CHE1[sym] OR DED[sym] OR apoptosis antagonizing transcription factor [GFN]) AND (MYC[sym] OR MRTL[sym] OR MYCC[sym] OR BHLHE39[sym] OR C-MYC[sym] OR MYC proto-oncogene [GFN])”. Note that there is no limit on the number of synonyms (ranging from 0 to 18) used in the query. Accessing our local dictionary is executed rapidly on the client side for this purpose. To estimate the cost of expanding the original query with synonyms and aliases, we compared the turnaround times of both queries on a TRRUST database. Figure 2 shows the boxplot of turnaround time for both queries. As can be seen, the time difference is trivial.

**Figure 2:**
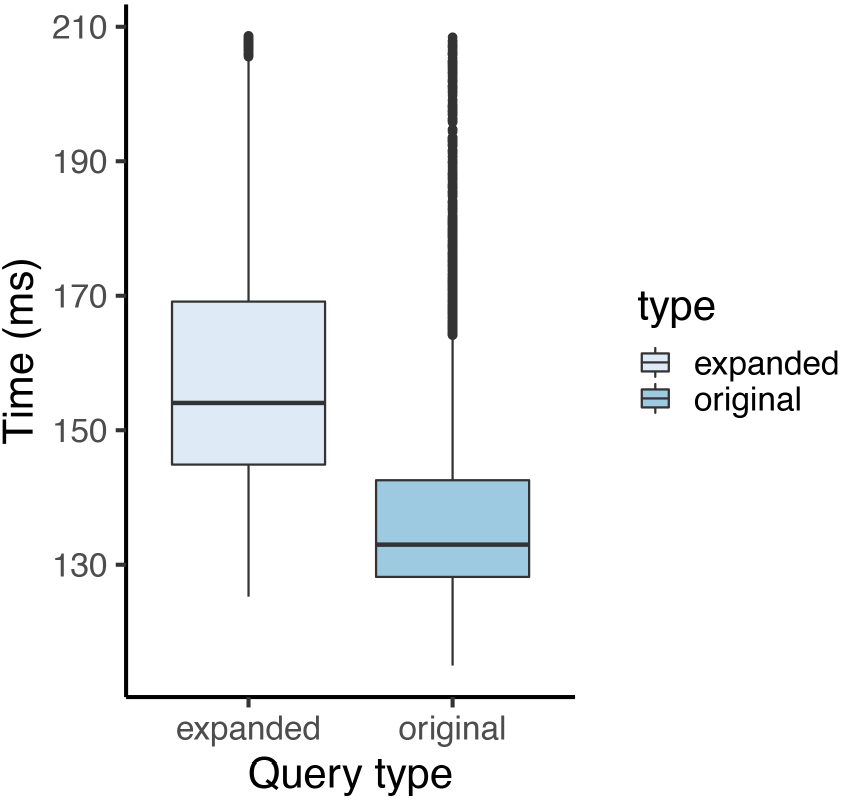
Box plot of turnaround time for expanded and original PubMed query on TRRUST database.

Finally, a dictionary-based approach was used to exclude irrelevant abstracts by scanning individual elements of them. We generated two sets of “causal regulatory events” including positive (activation) and negative (repression) events to purify the final abstracts. We applied a filter on retrieved abstracts and included only those abstracts which contain at least one regulatory event as presented in Table 1. Each category contains more than 50 verbs and their inflections. For example, the AATF-MYC query outlined above, resulted in 4 relevant abstracts (PMIDs: 20549547, 17006618, 17006618, 20924650).

**Table 1.** Regulatory events categories.

| Category | #Events | Examples |
| --- | --- | --- |
| Positive | 500 | increase, induce, activate, enhance |
| Negative | 511 | reduce, decrease, suppress, block |

### 3.3. Gene and regulatory entity recognition

The next step in the pipeline is to identify biological entities within the abstracts. Figure 3 shows the NER module of our system. Two external state-of-the-art NER systems were utilized to annotate the retrieved abstracts with an accurate and complete list of biological entities. The first system is PubTator [41], a web-based system for assisting biocuration. PubTator utilizes a HTTP REST interface, equipped with multiple state-of-the-art text mining algorithms to run query. Using this system, we queried the abstract PMIDs from IR module to PubTator interface and obtained entity annotations in a JSON encoded text. Additionally, we utilized BeCAS [42] (another online NER tool) to improve the coverage of the entities. BeCAS, like PubTator, provides a RESTful API for biomedical name identification. It can run queries directly on provided text or PMIDs and returns associated annotations as an XML document. Although both systems provide high consistent annotations for gene entities, we put more priority on PubTator when there is incompatible results for a particular entity.

**Figure 3:**
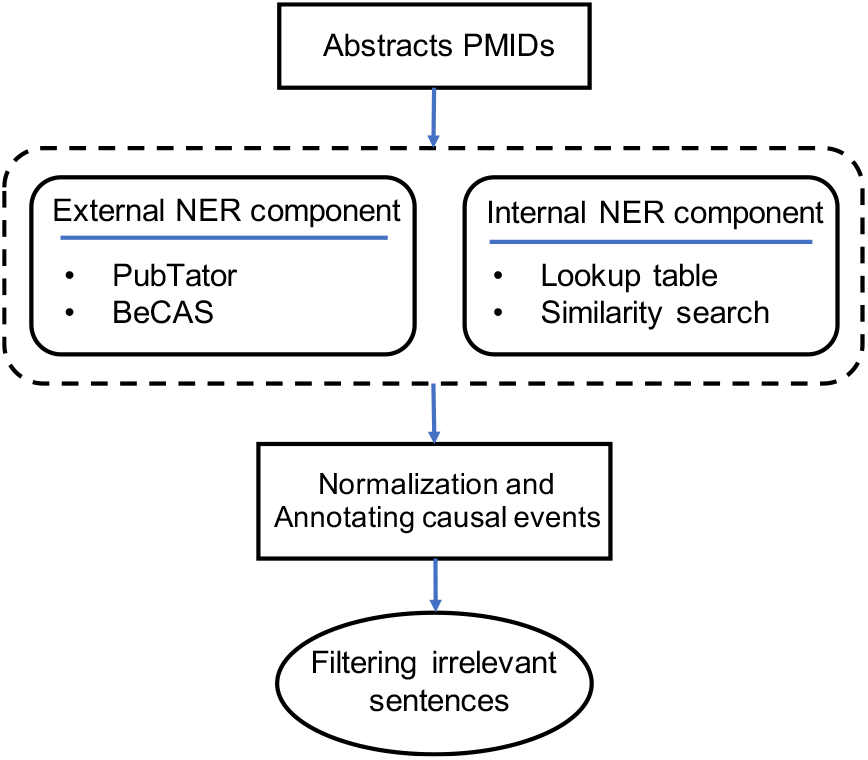
The gene entity and regulatory event recognition workflow. Each PubMed ID retrieved by IR component are submitted to the external NER tools (PubTator and BeCAS) for annotating genes in the abstracts. It follows complementary annotations using our internal NER component including a lookup table for covering acronyms, and a similarity search to identify lexical variations for gene names.

To further enhance the NER module, we implemented and added an additional NER component as follows. Abstracts were normalized to uppercase format and searched for gene acronyms using a manually-curated lookup table [43]. This table includes long term / short term pair association to recognize entities, which were missed by the external NER tools. For instance, AR is a short term for “Androgen Receptor” and was only detected as an entity (transcription factor) using this lookup table. Furthermore, we utilized a name similarity metric to identify strings with lexical variations such as whitespace and punctuations. For instance, “IL-12” and “IL12” are two lexical variations of “Interleukin 12”. The former version was not identified by the External NER systems. In our implementation, we set the entity detection threshold based on Jaro similarity [44] of 0.9 or larger between the query entity and the string in the abstract.

Finally, we normalized the annotated word or a group of words corresponding to a gene to their HGNC symbol for simplification of downstream analysis. Regulatory events were also annotated using our expert-generated categories (Table 1). Figure 4 illustrates the normalization of gene names and annotation of regulatory events. Importantly, in order to reduce noises, sentences that contained no regulatory event were excluded from further analysis. We used the remaining sentences from all citations to extract relation between the TF and target gene.

**Figure 4:**
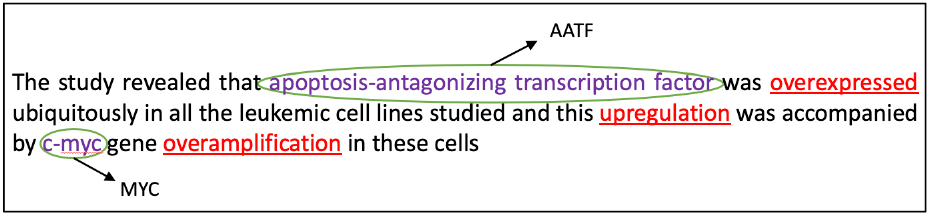
An example of gene entity normalization and regulatory events annotation. All of the words or group of words associated to target entities (purple color) are normalized to their HGNC symbol for simplification. Causal regulatory events also are annotated according to their categories, and sentences with no regulatory event are excluded for further consideration.

### 3.4. Extracting mode of regulation

Figure 5 illusterates the steps of the relation extraction workflow of our system. For each causal interaction, its annotated sentences from NER module were submitted to the Stanford dependency parser [45] and a dependency parse tree was generated. Dependency trees extracted from different sentences were merged into a single large graph. The merging process is straightforward; each dependency relation includes one head word/node and one dependent word/node. Nodes from different dependency relations representing the same word were merged together. PMID was recorded for each edge in the parse tree to indicate its source of evidence. Furthermore, each edge in the parse tree was assigned weight which is the number of occurrences of dependency relations with respect to all of the evidence sentences. The rational for using this weighted parse tree is that it can be used to identify long-range dependency relations across sentence boundaries that would otherwise be missed. Absolute frequency of a dependency relation obtained from the merging step can somewhat reflect the semantic relation of the head word and the dependent word.

**Figure 5:**
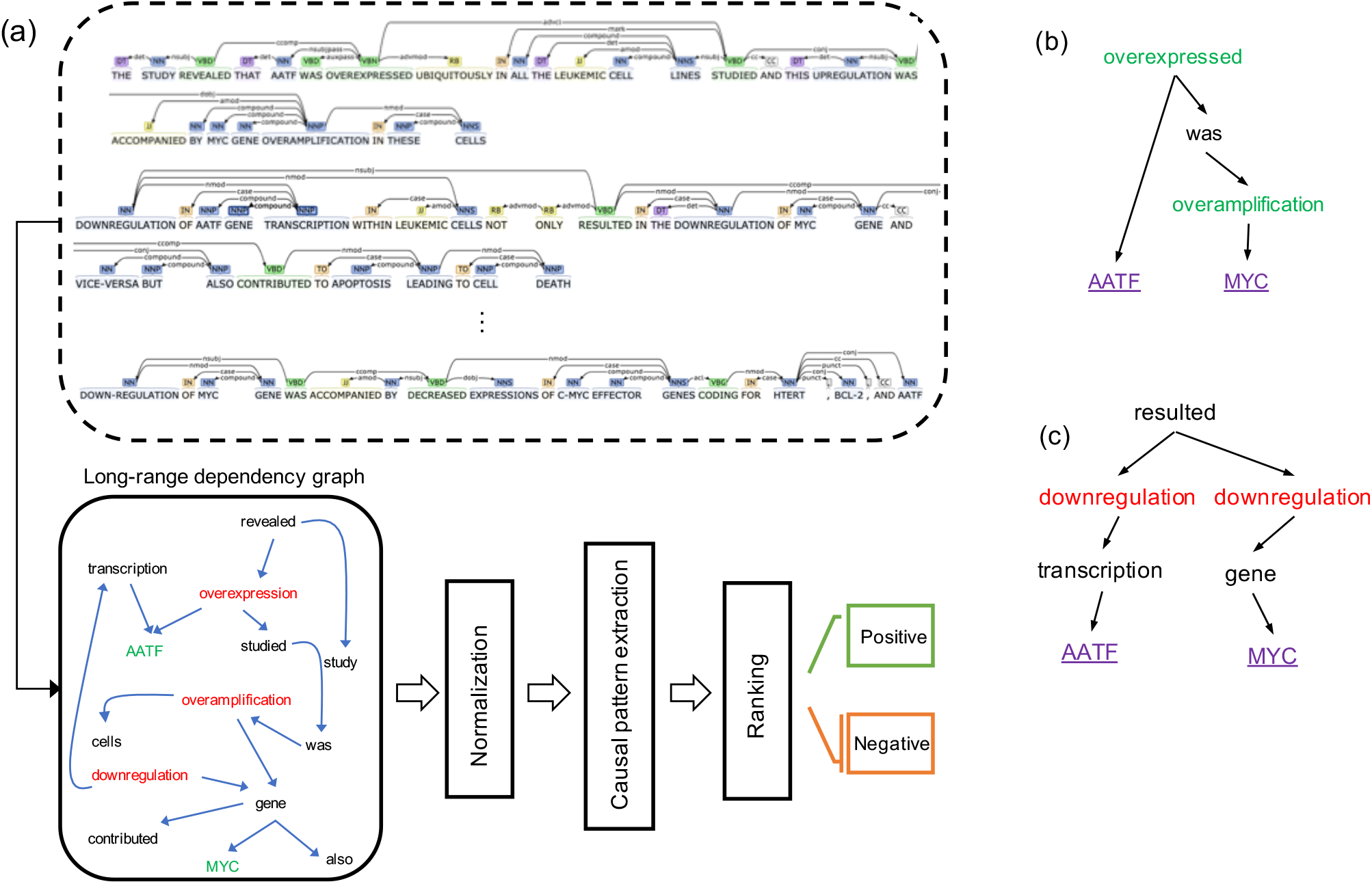
Relation extraction workflow extraction workflow. Panel (a) shows the construction of a long-range dependency graph by merging all of dependency trees corresponding to the evidence sentences. The weights of the graph reflect the number of occurrences of dependency relations. Candidate regulatory signs are identified using common subtrees with at least one regulatory event in the graph. Finally, a sign of regulation is assigned to the query interaction through the ranking task. panel (b) shows an example for simple rule (effector-relation-effectee) in which the RE system can assign a positive sign to this candidate pattern. In panel (c), we can see the impact of the negation rule to extract accurate sign to this pattern. Two paths from root to query entities contain negative regulatory events which carries an activation/positive sign for the pattern.

RE module creates candidate relations by extracting subtrees with common ancestors connecting the pair of query genes as leaves. These subtrees must contain at least one causal event describing the candidate relation between the given pair of genes. Subtrees were extracted by applying a Depth First Search along with a Boolean visited array to avoid possible loops. Nodes with two paths to the entities were considered as a root of the subtree. Next, we utilized a rule based approach to describe relations using three commonly used language constructs [29]. The first rule is effectorrelation-effectee (e.g. A activates B). The second rule is relation-of-effectee-by-effector (e.g. Activation of A by B). These rules were applied to both paths from root to query entities to identify their regulatory dependency. Figures 5b and 5c illustrate the regulatory relation extraction using these rules. Some sentences in the literature have complex structures, which cannot be captured by these language constructs. To address this, we incorporated a negation rule to increase the performance of the RE system. For example, consider the following sentence: “LMP1 suppresses the transcriptional repressor ATF3, possibly leading to the TGF-induced ID1 upregulation” [46]. In the first pass the system assigns a positive mode to the interaction between ATF3 and ID1. However, there is a negative interaction between the TF and target gene. The negation rule considers the negative event “suppresses” related to ATF3 and switch the positive mode to negative. Figure 5c shows a subtree reflecting the negation rule.

We then apply the rules to every subtree to extract the mode of regulation between the query genes. The weights of the graph encode repetition of regulatory relations across sentences and abstracts. we considered the weights when there was more than one regulatory event associated with the target gene. In this case, an event with higher weight was selected for ranking the subtree. We also considered distance of events to the target gene when the weights in the subtree were equal. The closest event to the target entity will take the highest priority for determining the interaction mode. Finally, we investigated regulatory mode in every candidate subtree and assigned a total mode of regulation to the interaction using a voting scheme. Algorithm 1 shows the algorithm used in ModEx to identify mode of regulation.

#### Algorithm 1

Algorithm for extraction of mode of regulation from evidence sentences.

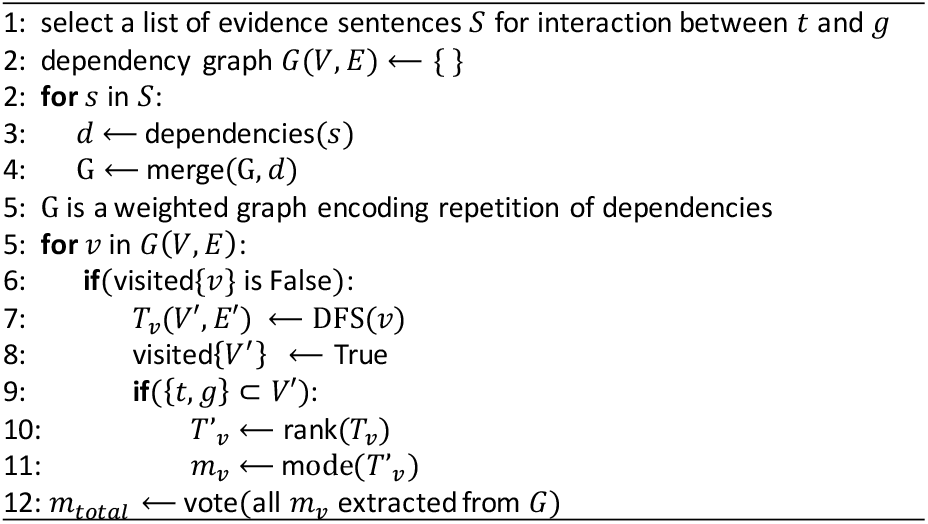

### 3.5. ModEx HTTP interface

We implemented an HTTP REST server for users to programmatically annotate gene regulatory networks using ModEx. Clients should make HTTP requests to the server with a particular format, specifying the query entities and optional MeSH term to annotate. The query has to be requested in the following format: TFEntrezID_TargetEntrezID_MeSHterm[optional]. For instance, a query to the server for AATF-MYC should be formatted as “/modex/26574_4609_humans”. Similar queries can be constructed by changing the Entrez ids. The server returns extracted annotation along with associated citations and sentences in XML format if any evidence exists. The turnaround time varies based on entities from one minute to a few minutes. For example, the server can be queried for the sample query AATF-MYC: https://watson.math.umb.edu/modex/26574_4609_humans.

## 4. Results

### 4.1. Impact of NER component

We tested the performance of different NER components of ModEx on gold-standard dataset provided by BioCreative V shared task 4 [47]. The dataset contains 7,555 manually annotated gene entities from 11,066 sentences from PubMed. For this experiment, we applied the NER system to sentences and compare the recognized gene entities with gold standard. We used two external NER components Pubtator and beCAS as standalone system, and evaluated their performance with our complementary internal NER functions (Section 3.3). We also report the performance of ensemble system including all of the components that we used in ModEx. Figure 6 shows the performance of various NER components of ModEx.

**Figure 6:**
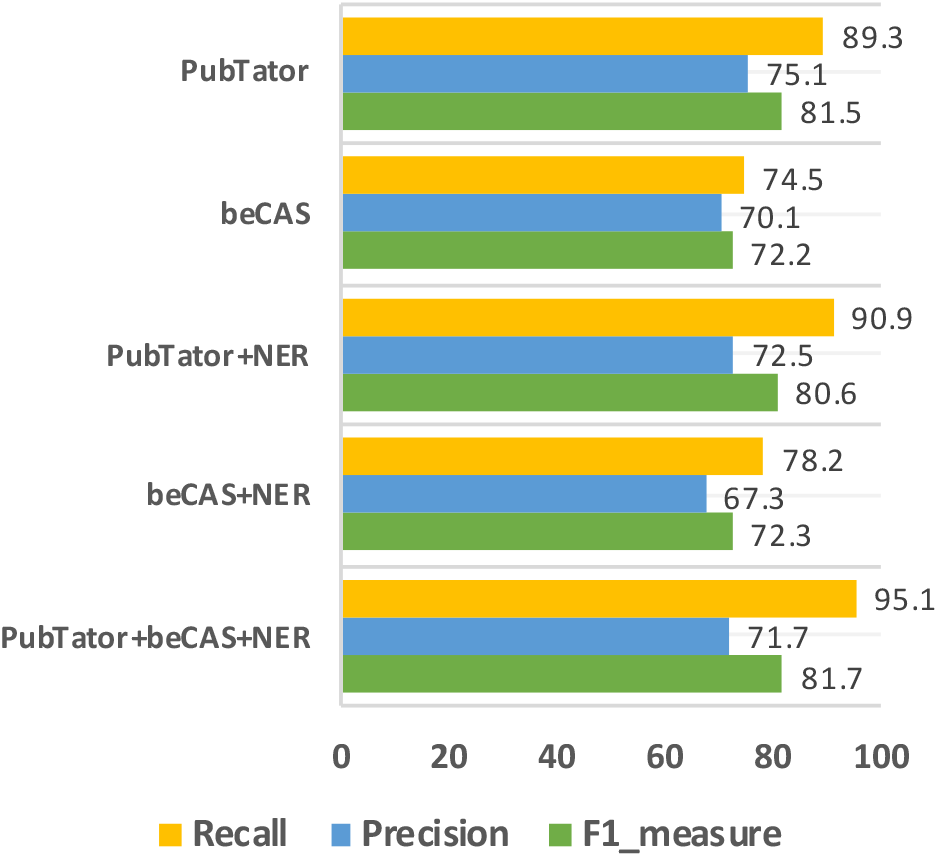
The performance result of five NER systems on the extraction of gene entities from gold standard.

The results indicate that all the systems achieved precision more than %67 for gene term extraction. Except for standalone BeCAS most other systems achieved very high recall. Note that we only investigated the performance of the systems on annotating gene entities. Although, our internal NER module compensated for the limitations of the external NER components, we did not observe a substantial increase in recall with marginal decrease in the precision of term extraction. Indeed, we achieved the highest performance in terms of F1-measure 81.7 using the ensemble NER system in ModEx.

### 4.2. Classification performance

We evaluated the performance of our method using the TRRUST database, a manually curated network or regulatory interaction with partial information on mode of regulation. TRRUST is a high-quality database and can be considered as gold standard for our benchmark. We applied our method to 5,066 regulatory interactions in TRRUST for which information on mode of regulation was available. As a benchmark, we developed an alternative pipeline using Integrated Network and Dynamical Reasoning Assembler (INDRA), a state-of-the-art text mining pipeline in biomedical domain, [48]. INDRA is an automated model assembly system interfacing with NLP systems and databases developed for molecular systems biology to collect knowledge and describe molecular mechanisms. In our approach, INDRA was used to assemble a reasoning model using causal statements extracted from literature through its submodules and methods. We configured INDRA to only extract, annotate and assemble a rule-based model to identify regulatory information from PubMed. The following steps were taken to assemble the pipeline. First, we used INDRA to mine PubMed using our expanded query for regulatory interactions. We then extracted all of the INDRA statement from returned abstracts using two standalone parsers, REACH [49] and TRIPS [50]. Finally, we incorporated all of the statements to assemble a reasoning model using PySB [51], a model assembler that implements a mathematical procedure to build a rule-based executable model. We identified inferred regulatory activity from the assembled model. Table 2 shows a summary of output of performing ModEx and INDRA on TRRUST database. INDRA identified PubMed abstracts corresponding to 4,942 of the annotated regulatory interactions in TRRUST, while ModEx extracted 4,225 abstracts due an additional filtering step based on regulatory events in the IR module (Section 3.2). ModEx and INDRA detected 4,225 and 3,093 regulatory activity (mode) respectively that are divided into activation and repression. We compared 3,077 interactions of intersection between the ModEx and INDRA results with the reported regulatory activities in TRRUST database. Figure 7 outlines the classification performance of ModEx and INDRA to identified mode of regulation. The result shows that our method outperforms INDRA with F1Measure 0.76 in prediction of mode of regulations.

**Table 2.** Summary statistics of performing ModEx and INDRA on TRRUST.

| ModEx |  |  | INDRA |  |  |
| --- | --- | --- | --- | --- | --- |
| #Abstracts | Extracted activity |  | #Abstracts | Extracted activity |  |
|  | 4,225 |  |  | 3,093 |  |
| 4,884 | Activation<br>2,659 | Repression<br>1,566 | 4,942 | Activation<br>2,173 | Repression<br>920 |

**Figure 7:**
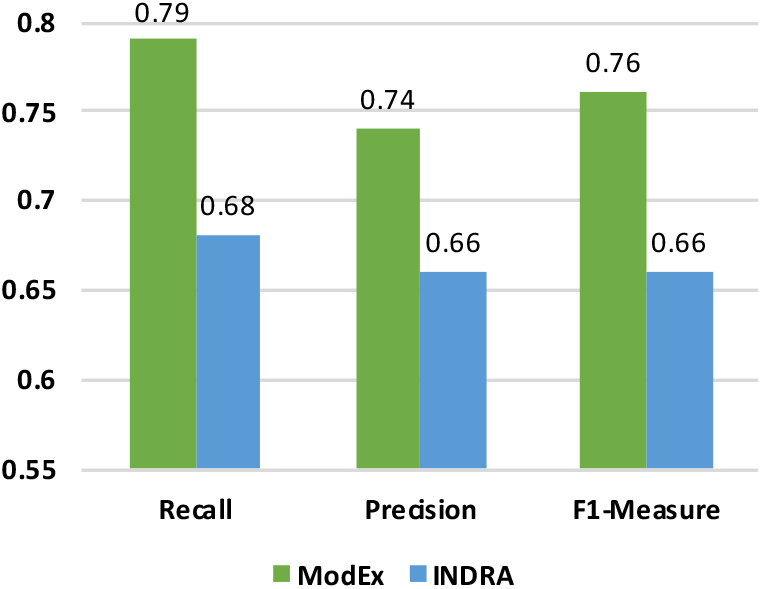
Classification results of ModEx and INDRA on TRRUST.

### 4.3. ChIP-Atlas analysis

We next sought to extract and annotate ChIP-seq derived TF-gene causal regulatory interactions from literature using our system. Such meta-data and evidence from literature can increase the confidence in the TF-gene interactions identified by ChIP-seq experiments and further shed light on the mechanism of interaction. Information on mode of regulation in particular can be helpful to enhance the accuracy of enrichment algorithms for regulatory pathway inference [55].

We applied ModEx to ChIP-seq interactions, with moderately stringency criteria, i.e., binding distance within 1k of the TSS and ChIP peak score > 950, resulting in 43,444 interactions. The system was able to detect and annotate 1,592 of interactions in PubMed database. Table 3 outlines the summary of output result on ChIP-Atlas.

**Table 3.** Summary statistics of performing ModEx on ChIP-Atlas.

| Overall | With evidence | With regulatory mode |  |
| --- | --- | --- | --- |
|  |  | 1,592 |  |
| 43,444 | 5,133 | Positive<br>1,421 | Negative<br>171 |

Some of the retrieved annotated ChIP-seq interactions also appear in the TRRUST database (69 total), indicating the low coverage of the TRRUST database. We compared the identified mode of regulations of ChIP-Seq interactions with the reported ones in the TRRUST database. Figure 8 summarizes the classification results. As can be seen the agreement is very high, indicating that our method can reliably identify and annotate ChIP interaction when they are reported in literature. Additionally, we compared our acquired evidence (PMIDs) by ModEx with citations reported in TRRUST. Our IR module was able to fetch the relevant evidence from PubMed database with accuracy 0.88.

**Figure 8:**
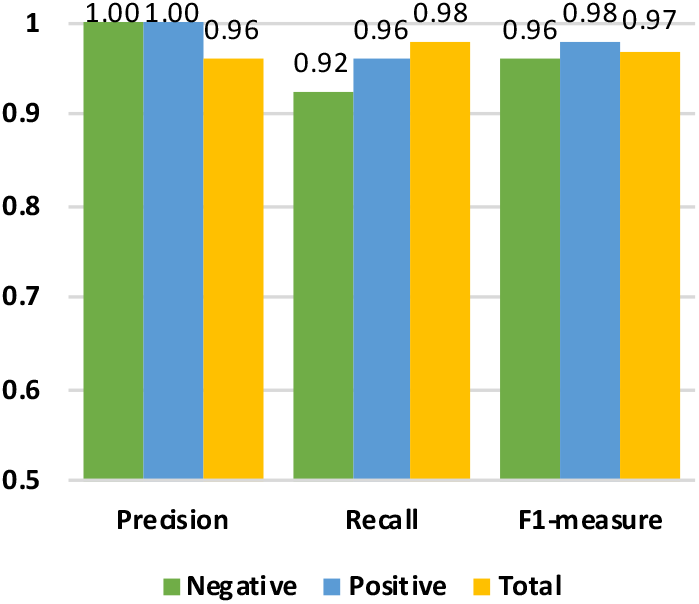
Classification results of ModEx on intersection on TRRUST and ChIP-Atlas.

### 4.4. Directional enrichment analysis

To demonstrate the utility of our annotated network, we used our network in conjunction with a directional enrichment analysis algorithm [52, 53] to identify drivers of differential expressed genes. We utilized quaternaryProd, a gene set enrichment algorithm that can take advantage of direction of regulation on causal biological interaction graphs to identify regulators of differential gene expression. The quaternaryProd algorithm takes a differential gene expression profile along with an annotated transcriptional regulatory network, such as TRRUST or our ChIP-Network as input and outputs a set of candidate active protein regulators. The algorithm performs a directional enrichment test based on the availability of information on the mode of regulation in the network. If no information on mode of regulation is available, the algorithm performs the Fisher’s exact test or the enrichment scoring (ES) statistic, which is the standard for gene set enrichment analysis. If the network if fully annotated, a generalization of the Fisher exact test (correctness p-value) is performed. For networks with both signed and unsigned edges, the algorithm performs Quaternary scoring (QS) statistic proposed by Fakhry et. al [52].

The Network inputted to quaternaryProd is assumed to encapsulate knowledge of TF-DNA interactions. The algorithm can use information on mode of regulation to more accurately identify putative protein regulators. The ability of the algorithm to identify regulators of differential gene expression relies heavily on the quality and the coverage of the regulatory network on which the queries are performed. If the network adequately encapsulates the interaction between TF and genes, the expectation is that the quaternaryProd algorithm should be able to recover the true cause of the modulated expression profile. To test the utility of our network, we used this algorithm along with differential expression profiles from controlled over-expression experiments used in the original study. The over-expression experiments consist of differential gene expression profile from a controlled in vitro E2F3 over expression [54] and c-Myc [54]. These over-expression experiments provide an ideal setting to test whether the network provides adequate and accurate information for the algorithm to recover the perturbed regulator or its closely related proteins. We inputted three networks into the algorithm (1) the original TRUSST network, (2) annotated TRUSST network, and (3) annotated TRRUST augmented with annotated ChIP-Atlas. By annotated TRRUST, we refer to the TRRUST network where interaction with no reported mode of regulation were annotated using our system. Differential gene expression analysis of these data sets resulted in 272, and 220 differentially expressed genes respectively. Table 4 outlines the top 10 regulators predicted by the algorithm on E2F3 differentially expressed genes sorted by the FDR corrected quaternary p-values of the scoring scheme (See Supp. File S1 for all predicted regulators with adj. p-value < 0.05). For the E2F3 experiment, E2F1 is returned as the top hypothesis regulator by the algorithm incorporating our annotated networks. E2F1 and E2F3 are close family members and have a very similar role as transcription factors that function to control the cell cycle and are similarly implicated in cancer [55]. It is interesting to note that original TRRUST database does not include enough information for algorithm to recover E2F1, however the signal strengthens when TRUSST is annotated with our system and a much more significant p-value is obtained when TRRUST is augmented with annotated ChIP-Atlas. This shows that annotating ChIP-seq data provides significant additional power to identify upstream regulators in conjunction with freely available causal networks.

**Table 4.** Directional enrichment analysis results on E2F1 expression signatures.

| TRRUST |  |  | Annotated TRRUST |  |  | Annotated TRRUST with ChIP-Atlas |  |  |
| --- | --- | --- | --- | --- | --- | --- | --- | --- |
| Name | Regulation | Adj. pval | Name | Regulation | Adj. pval | Name | Regulation | Adj. pval |
| REL | Down | 1.5e-3 | E2F1 | Up | 1.2e-5 | E2F1 | Up | 1.2e-7 |
| PROX1 | Up | 2.9e-3 | PROX1 | Up | 2.1e-3 | PROX1 | Up | 2.1e-3 |
| SUGP1 | Up | 3.2e-3 | SUGP1 | Down | 2.4e-3 | SUGP1 | Down | 2.4e-3 |
| NFIL3 | Up | 6.3e-3 | RELA | Down | 3.5e-3 | RELA | Down | 3.5e-3 |
| TFDP1 | Up | 7e-3 | TFDP1 | Up | 3.9e-3 | TFDP1 | Up | 3.9e-3 |
| ARNTL | Down | 7.5e-e3 | STAT5 | Down | 3.9e-3 | CLOCK | Up | 3.9e-3 |
| NPAS2 | Up | 1.2e-2 | ARNT | Up | 5.7e-3 | ARNT | Up | 5.7e-3 |
| CLOCK | Down | 1.2e-2 | SNW1 | Up | 6.6e-3 | STAT5 | Down | 6.0e-3 |
| AHR | Up | 1.2e-2 | IKZF1 | Up | 8.9e-3 | SNW1 | Up | 6.6e-3 |
| HDAC1 | Down | 1.4e-2 | GATA4 | Down | 1.4e-2 | IKZF1 | Up | 8.9e-3 |

Application of the method to c-Myc differential expression profile shows the similar pattern. The annotated TRRUST with ChIP-Atlas recovered MAX as one of the predicted regulators which were not identified using TRRUST and annotated TRRUST networks. (See Supp. File 2 for all predicted regulators with adj. p-value < 0.05). It has been demonstrated that oncogenic activity of c-Myc requires dimerization with MAX [56].

**Table 5.**
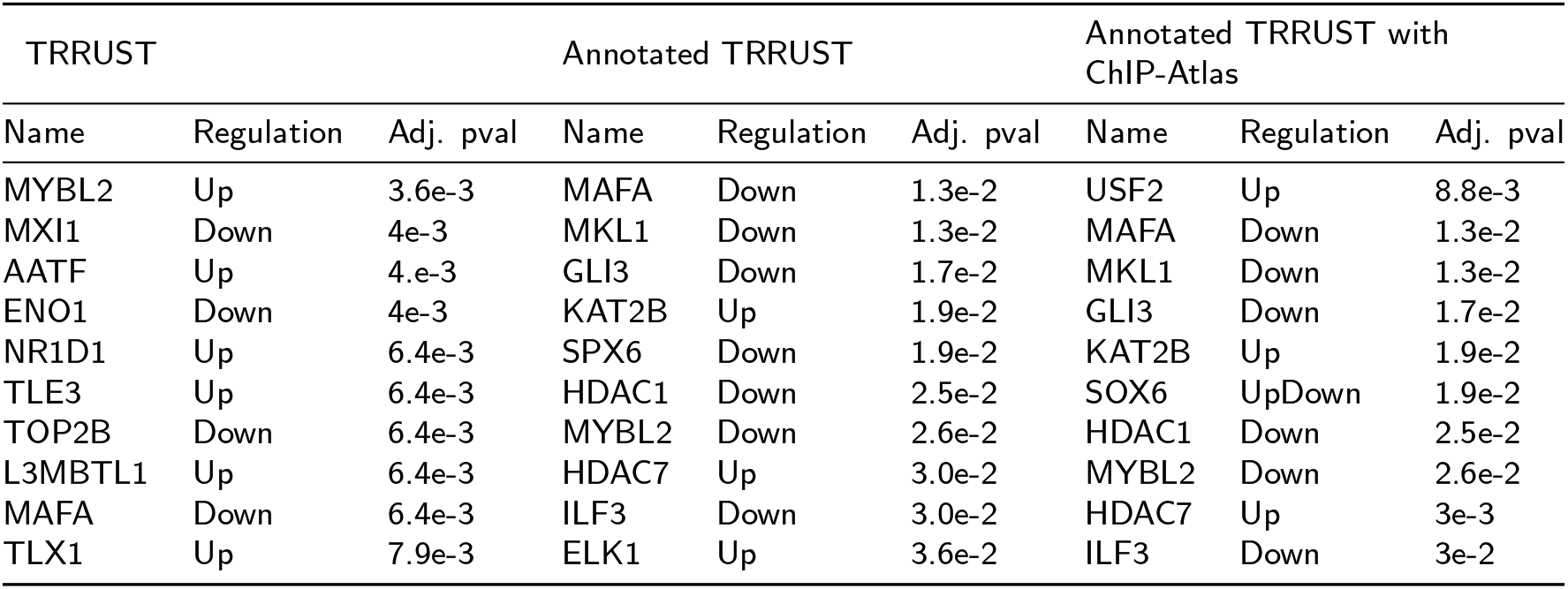
Directional enrichment analysis results on c-Myc expression signatures.

## 5. Conclusion

In this work we presented a fully automated text-mining system to extract and annotate causal regulatory interaction between transcription factors and genes from the biomedical literature. As a starting point, our method uses putative TF-gene interactions derived from high-throughput ChIP-seq or other experiments and seeks to collect evidence and metadata in the biomedical literature to support the interaction. It should be noted that annotating a priori known interactions differs significantly in scope and complexity from general text-mining approaches for biomedical relation extraction. The later attempts to extract the causal relation from biomedical text directly, without prior knowledge of the entities and the interaction, whereas in our method the relation is know from biological experiments and curated databases a priori, thereby reducing the complexity significantly. This approach bridges the gap between data-driven methods and text-mining methods for constructing causal transcriptional gene regulatory networks and overcomes some of the drawbacks of either approach. With the rapid increase in high-throughput experiments and biomedical literature, hybrid method such as the one proposed can make a significant impact in biological knowledge retrieval.

We used a gold-standard manually curated dataset and demonstrated that our approach can reliably identify the relevant literature and extract the correct interaction and metadata. We applied our method to high-throughput ChIP-seq data and provided literature support for 1,500 interactions. Our annotated ChIP-derived transcriptional regulatory interaction can be used in conjunction with directional enrichment methods that aim to identify regulators of differential gene expression. Moreover, we use our system to annotate the interactions in the TRRUST database for which more of regulation is not reported. Our system can also be used as a tool to mine the literature for investigate interactions in newly performed ChIP-seq experiments, where researchers are interested to investigate a specific interaction between a protein and a gene. To facilitate usage, we implemented an HTTP REST server for users to programmatically annotate gene regulatory networks using ModEx available via: https://watson.math.umb.edu/modex/[type_query] (See section 3.5). The annotated ChIP-network as well as annotated TRRUST can be obtained from: https://doi.org/10.6084/m9.figshare.8251502.v1

